# The evolutionary demise of a social interaction: social partners differ in the rate at which interacting phenotypes are lost

**DOI:** 10.1101/2022.04.05.486946

**Authors:** Eleanor K. Bladon, Sonia Pascoal, Nancy Bird, Rahia Mashoodh, Rebecca M. Kilner

**Affiliations:** University of Cambridge; University College London

## Abstract

Phenotypic plasticity enables animals to adjust their behaviour flexibly to their social environment – sometimes through the expression of adaptive traits that have not been exhibited for several generations. We investigated how long social adaptations can usefully persist when they are not routinely expressed, by using experimental evolution to document the loss of social traits associated with the supply and demand of parental care. We allowed populations of burying beetles *Nicrophorus vespilloides* to evolve in two social environments for 48 generations in the lab. In ‘Full Care’ populations, traits associated with the supply and demand of parental care were expressed at every generation, whereas in ‘No Care’ populations we prevented expression of these traits experimentally. We then revived trait expression in the No Care populations at generations 24, 43 and 48 by allowing parents to supply post-hatching care, and compared these social traits with those expressed by the Full Care populations. We found that offspring demands for care decayed in the No Care populations more rapidly than a parent’s capacity to supply care. Furthermore, male care decayed before female care. We suggest that this reflects differences in the strength of selection for the expression of alternative traits in offspring, males and females, which can enhance fitness when post-hatching care is disrupted.

**Impact Summary:** Social interactions between animals are suggested to be increasingly vulnerable to breakdown in our changing world. Our experiments offer a rare insight into what happens next, by assessing in real time the durability of social behaviours that are no longer routinely expressed. Our results also have implications for conservation captive breeding programmes where compensatory husbandry techniques prevent trait expression and so could inadvertently induce rapid, irreversible trait loss.

We investigated how long it took populations to lose the ability to express appropriate social behaviour when they had been prevented from doing so for many generations. We did this by evolving replicate populations of burying beetles *Nicrophorus vespilloides* in the laboratory for 48 generations. The burying beetle is a common insect that is well-known for caring for its larvae, although larvae can survive in the lab without any care at all. In two populations (“Full Care”), we allowed parents and offspring to interact during the supply of post-hatching care, as usual. In two other populations (“No Care”), parents were removed before offspring hatched and so could not interact socially with their young.

Over the course of 48 generations of experimental evolution, we periodically revived social interactions between parents and offspring in the No Care populations. We assessed the extent to which larval begging behaviours, and parental care behaviours, had decayed by comparing their expression with those in the Full Care populations. We found that larval begging behaviour eroded rapidly in No Care populations, and more rapidly than the supply of care by parents. Furthermore, paternal care decayed to a greater extent than maternal care (which was largely unchanged relative to its expression in the Full Care populations). We suggest that these differences could be due to differences in the strength of selection on each family member for alternative traits to enhance fitness.

## Introduction

Phenotypic plasticity enables animals to flexibly, and rapidly, adjust their behaviour according to the environment in which they live – sometimes through the expression of adaptive traits that have not been exhibited for several generations (Davies, 2000; Lahti, 2006). The ability to revive ‘ghosts of adaptations past’ could prove beneficial for populations living in a changing world (Lahti, 2006; Robert and Sorci, 1999; Rothstein, 2001). For this reason, it is important to understand how long such adaptive traits can persist at the population level if they are no longer routinely expressed. This is especially true for behavioural traits, which have been relatively under-studied in this context (Rayner et al., 2022).

The capacity for any unexpressed trait to persist is thought to be due to a combination of adaptive and neutral processes (Lahti et al., 2009, Rayner et al. 2022). Adaptive processes are involved because selection will act against the mechanisms that enable an unexpressed trait to be environmentally induced if there are associated costs with trait plasticity (DeWitt et al., 1998; Snell-Rood et al., 2018). The costs could be outweighed by any fitness benefits accrued through the occasional expression of this trait. If there are no net costs, then the trait might persist – vestigially and non-adaptively (Rayner et al., 2022). However, the greater the net fitness costs, the more rapidly we would expect the mechanism for plasticity to be lost. There are then two possibilities – either there is genetic assimilation and the trait is constitutively expressed, independent of environmental conditions (DeWitt et al., 1998; Pigliucci et al., 2006; Robinson and Dukas, 1999; Scheiner and Levis, 2021; Snell-Rood et al., 2009; Waddington, 1953), or trait expression ceases altogether and the trait is lost (Lahti, 2006; Rayner et al., 2022). The latter possibility is the focus of our work here.

Neutral processes can also explain how previously plastic traits are lost over time – especially in the longer-term. If a trait is unexpressed for many generations, then it is not exposed to selection and deleterious mutations can potentially accumulate in the underlying genes (Snell-Rood et al., 2009). Eventually, this too could cause the trait to be lost forever (Lahti et al., 2009). Nevertheless, the initial stages of trait loss have seldom been observed directly, partly because these steps involve relatively subtle nuanced change rather than wholesale disruption (Ellers et al., 2012).

In this study, we used experimental evolution to document the loss of unexpressed behavioural traits in real time. Our work focused on plastic social traits that are induced through interactions with a conspecific, namely the demand and supply of parental care. It has recently been suggested that social interactions like this are likely to be disrupted by broader environmental change, though the evolutionary implications are still largely unknown (Bailey and Moore, 2018). Theoretical work generally suggests that social interactions can enhance the pace of evolutionary change through indirect genetic effects, not only by increasing the pace at which new socially expressed traits spread through the population but also by potentially accelerating their demise, by disrupting or preventing the expression of traits in a social partner (McGlothlin et al., 2010; Moore et al., 1997; Wolf et al., 1998). We mimicked this type of social disruption by experimentally preventing the expression of post-hatching parental care in the burying beetle *Nicrophorus vespilloides* in some populations (“No Care”) for 48 generations, whilst allowing its continued expression in other populations (“Full Care”). After 24, 43 and 48 generations of experimental evolution, we revived interactions between parents and offspring in the No Care populations and compared the expression of the parental and offspring traits involved with the Full Care populations, to measure any changes in the frequency of their expression at the population level and the subsequent fitness effects.

We predicted (1) that by generation 24, and even more so by generation 43, we would see a decay in parental and/or larval traits associated with the supply and demand of parental care. We assessed this indirectly by measuring larval mass, since it increases with the supply of parental care (Bladon et al., 2020; Pascoal et al., 2018; Steiger, 2013). We isolated parental and larval contributions to larval mass independently, by cross-fostering larvae between Full Care and No Care populations. We predicted that if No Care larvae had lost the capacity to demand care, then they should be less effective than Full Care larvae at gaining mass when raised by Full Care parents. Conversely, if No Care parents had lost the capacity to supply care, then Full Care larvae should gain less mass when raised by No Care parents than when raised by Full Care parents.

Additionally, we quantified the supply and demand for parental care directly. We predicted (2) that if No Care parents had lost the ability to supply care then the duration of their care should be shorter than shown by Full Care parents (Eggert et al., 1998). We also predicted (3) that if No Care larvae had lost the ability to demand care then they should be less inclined to solicit care from parents (Smiseth and Parker, 2008; Smiseth et al., 2010).

## Methods

### *Natural History of Burying Beetles* Nicrophorus vespilloides

Burying beetles show elaborate parental care that is highly variable in its duration (Jarrett et al. 2018b). Male and female beetles can detect small vertebrate carrion from a distance of several kilometres with their highly sensitive antennae (Kalinová et al., 2009). A pair converts the dead body into a nest for their young by tearing off the fur or feathers, covering the flesh with antimicrobial anal and oral exudates, rolling it into a ball and burying it underground (Cotter and Kilner, 2010). Meanwhile, the pair mates and the female lays her eggs in the soil around the nest. The larvae crawl to the nest after hatching (Müller and Eggert, 1989). Both parents tend the larvae by defending them from intruders and supplying fluids through oral trophallaxis (Eggert et al., 1998). Larvae also self-feed (Smiseth et al., 2003) and can survive (at least in the laboratory) without any post-hatching care, though they attain a lower mass at dispersal and have a poorer chance of surviving to independence (Eggert et al., 1998). Larval mass is a key correlate of fitness because larval mass predicts adult size (Jarrett et al., 2017) which, in turn, predicts fecundity in both males and females (Bladon et al., 2020; Pascoal et al., 2018).

### Burying beetle husbandry in the laboratory

For all breeding experiments, each pair of sexually mature male and female beetles was bred by placing them in a plastic breeding box (17 × 12 × 6 cm) with damp soil (John Innes Compost) and a 10-15 g mouse carcass on which to breed. Larvae were counted and weighed 8 days after pairing and placed in plastic pupation boxes (10 × 10 × 2 cm), filled with damp peat. Sexually immature adults were eclosed approximately 21 days later and housed in an individual box (12 × 8 × 2 cm). Adults were fed twice a week (beef mince) until breeding, which took place 15 days post-eclosion. Adults and pupating larvae were kept on a 16L: 8D hour light cycle at 21°C.

### Experimental Evolution

The *N. vespilloides* populations described in this study were part of a long-term experimental evolution project that investigated how populations of burying beetles adapt to the loss of parental care. It focused on four experimental populations: Full Care (FC; x2 replicates) and No Care (NC; x2 replicates). Their establishment and husbandry have been described in detail before (Duarte et al., 2021; Jarrett et al., 2018a, 2018b; Rebar et al., 2020; Schrader et al., 2015a, 2015b, 2017). Briefly, these populations were established in 2014 with wild-caught beetles (trapped under permit) from four woodland sites across Cambridgeshire, UK (Byron’s Pool, Gamlingay Woods, Waresley Woods and Overhall Grove). The NC populations were routinely prevented from supplying any post-hatching care, through the removal of adults at 53 h post-pairing, when the carrion nest was complete but before the larvae had hatched. Whereas in the FC populations adults were allowed to stay with their larvae throughout development and provide care. This procedure was repeated at every generation. Each type of experimental population was run in a separate block (FC1/NC1 and FC2/NC2) with breeding staggered between blocks by 7 days. We used these populations to assess the level of parental care populations provided towards offspring and the amount of care solicited in offspring at generations 24, 43 and 48.

We have previously described the divergent adaptive evolution of these experimental populations in response to the loss of care. Although the No Care populations initially showed higher rates of larval mortality, they swiftly adapted (Schrader et al., 2015b). After 23 generations of experimental evolution, larvae had a similar chance of survival and attained a similar mass at dispersal in each type of population (Rebar et al., 2020; Schrader et al., 2017). Over the same time frame, No Care larvae evolved to hatch more synchronously (Jarrett et al., 2017), to have disproportionately larger mandibles (Jarrett et al., 2018a), and to be more cooperative with their siblings (Jarrett et al., 2018b; Rebar et al., 2020). No Care parents evolved to frontload parental care, making a rounder carrion nest more quickly than Full Care parents, which promoted larval survival in the absence of post-hatching care (Duarte et al., 2021).

#### Experiment 1: Assessing the duration of care, and partitioning larval and parental contributions to brood mass

We assayed the propensity for parents to supply care in the evolving populations after 24 and 43 generations of experimental evolution. Whilst they are providing parental care, parents typically stay in close proximity to the carcass. We measured the duration of care exhibited by NC and FC parents by breeding them in a box with a plastic partition that allowed parents to desert the nest after parental care was completed (Figure S1; (de Gasperin et al., 2015)). We therefore used time of desertion as a proxy for when parents had stopped supplying parental care.

Each breeding pair was placed in the larger compartment (Figure S1), along with an 8-12 g carcass. At 53 h post-pairing, we cross-fostered parents between boxes to create families that either stayed with their own nest or were translocated to a foster nest of the same (control) or different (cross-fostered) type of experimental population (i.e. No Care or Full Care) (see Figure S2). This design enabled us to partition the contributions of parents versus offspring to brood mass at dispersal. The transfers within experimental populations enabled us to control for any effects due to the cross-fostering procedure itself. Following the translocation of some pairs and nests, adults remained in the boxes until they entered the escape chamber or the experiment ended (8 days after pairing). Thus, in all treatments, parents were able to supply post-hatching care. From breeding to dispersal, we checked the escape chamber every 4 hours between 8am and 8pm (i.e. 64-192 h after pairing in generation 24; 56-192 h after pairing in generation 43). At dispersal (8 days post-pairing), we removed the remaining parents, counted the number of surviving larvae and weighed the brood.

#### Experiment 2: Assessing larval begging behaviour

We quantified the frequency of larval begging behaviour using previously described methods (Smiseth and Parker, 2008; Smiseth et al., 2010). At generation 48, we created Full Care pairs and No Care pairs of sexually mature, unrelated, virgin beetles and put them in a standard size breeding box (17 × 12 × 6 cm) with 300 ml soil and a 10-13 g mouse carcass. At 53 h after pairing, we removed the male and transferred the female and her carrion nest into a new standard size breeding box (also containing 300 ml soil). We checked the original breeding box for freshly-hatched larvae 24 h later. Larvae were pooled by condition and distributed to create four conditions: 1) Full Care female with 15 Full Care pooled larvae, 2) Full Care female with 15 No Care pooled larvae, 3) No Care female with 15 Full Care pooled larvae and 4) No Care female with 15 No Care pooled larvae. This design ensured that begging was not specific to parent of origin. Larvae were placed directly on the focal female’s carrion nest. Females were included in the experiment only if their original eggs hatched successfully to prevent cannibalism of larvae that appear prior to their own eggs hatching (Müller and Eggert, 1990; Smiseth and Parker, 2008).

Twenty-four hours after establishing the experimental broods, when the larvae were second instar and had reached peak begging activity (Smiseth et al., 2003), we removed the females from their broods, and placed them in labelled containers in a −20 °C freezer for 30 min to euthanise them. Meanwhile, all surviving larvae were removed from their brood ball and placed on labelled pieces of damp paper towel for 25 min prior to the start of the experiment to increase the solicitation of care (T Ratz. *pers.comm.*). After removal from the freezer, each female was thawed for 5 min and mounted on a pin at the centre of a plastic box (11 × 17 × 4.5 cm) lined with damp paper towel, mimicking the stance of a parent regurgitating food (Mäenpää and Smiseth, 2017).

The focal female’s experimental brood was then added to the container, with individual larvae placed haphazardly within it, and left for 5 min to acclimatise. We used instantaneous scan sampling to detect begging activity (Mäenpää and Smiseth, 2017; Smiseth and Parker, 2008; Smiseth et al., 2010), recording larval activity every minute for a 10 min period. Activity was classified as either 1) associating with the parent (a larva was within one-pronotum’s width of the female); or 2) begging (a larva was rearing up and touching the parent with its legs); or 3) neither.

#### Statistical analyses

All statistical tests were conducted in R version 3.6.1 (R Core Team, 2019). Data handling and visualisation were carried out using base R and the ‘tidyverse’ suite of R packages (Wickham et al., 2019). A stepwise deletion method using F tests, implemented in the base ‘statistics’ package in R, was used to determine significance of each term and remove non-significant terms sequentially (Crawley, 2007).

### Experiment 1

Prediction 1: By generation 24, and even more so by generation 43, we should see a decay in parental and/or larval traits associated with the supply and demand of parental care, as inferred through parental and larval contributions to larval mass.

We analysed generation 24 and 43 separately using a linear model with a Gaussian error structure, implemented by base R regression functions. The dependent variable was brood mass at dispersal (g) and the predictor variables included in the maximal model were brood size, carcass mass (g), current parents’ experimental population (No Care or Full Care), larval experimental population (No Care or Full Care), whether the parents had been transferred between carcasses and eggs or not, male leaving time, female leaving time and experimental block (1 or 2) and the interaction between current parents’ experimental population and larval experimental population.

Prediction 2: If No Care parents had lost the ability to supply care then the duration of their care should be shorter than shown by Full Care parents.

We first conducted an ANOVA (using the base R ‘statistics package) to determine whether there was a significant difference in male and female leaving time across generations. As there was a significant difference, male and female parents were then analysed separately in subsequent analyses. We analysed the parental leaving time data using semi-parametric Cox’s proportional models for interval censored data (using the icenReg R package (Anderson-Bergman, 2017)). In the male maximal models we included parental experimental population (No Care or Full Care), focal offspring experimental population of origin (No Care or Full Care), whether the parents had been transferred or not, number of larvae, mass of mouse carcass and experimental block as fixed effects. In the female maximal models, the same terms were included but male leaving time was also included to determine whether females were more likely to leave earlier once their partner left. Female leaving time was not included in the male model because there were so few instances of females leaving before their partner (n = 9/153 in generation 24; 13/214 in generation 43).

### Experiment 2

Prediction 3: If No Care larvae had lost the ability to demand care then they should be less inclined to solicit care from parents.

To determine whether there were differences in begging duration between larvae originating from the Full Care and No Care experimental populations, we first ran a quasi-binomial generalised linear models (GLM) with begging duration as the response variable. In a second model, we tested for differences in any behaviour when larvae interacted with the (dead) parent by combining begging duration with time spent associating with the parent. In both models, the predictor variables included in the maximal model were larval experimental population (No Care or Full Care), foster female’s experimental population (No Care or Full Care), number of surviving larvae in the brood, experimental block and the interaction between larval experimental population and foster female’s experimental population.

### Results

#### Experiment 1

Prediction 1: By generation 24, and even more so by generation 43, we should see a decay in parental and/or larval traits associated with the supply and demand of parental care, as inferred through parental and larval contributions to larval mass.

In generation 24, larval experimental population accounted for more variation in brood mass at dispersal than did parental experimental population. Broods of larvae from the No Care experimental populations were significantly lighter than those from the Full Care experimental populations (linear regression: *F*_1,147_ = 30.84, *p* = <0.001, Figure 1A), regardless of the experimental population of origin of the parents that cared for them (linear regression: *F*_1,144_ = 0.870, *p* = 0.353).

**Figure 1.**
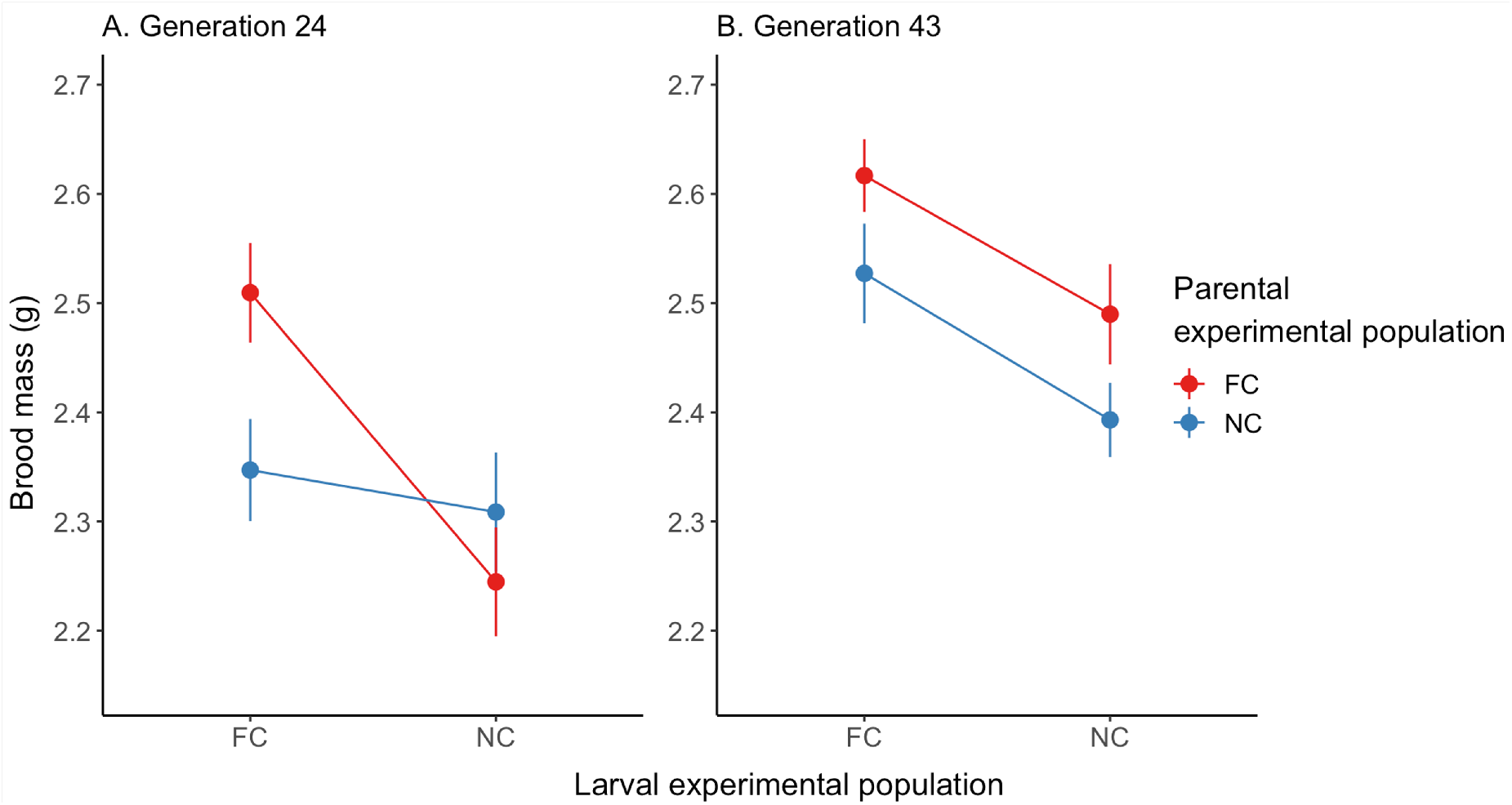
The effect of parental experimental population and larval experimental population on brood mass at dispersal in A) generation 24 (n = 153) and B) generation 43 (n = 214). Points represent predicted values from the minimal model containing both parental experimental population and larval experimental population terms. In A) larval experimental population explains significant variation in brood mass at dispersal, but neither parental experimental population, nor the interaction between the two, are significant, and in B) both larval and parental experimental population explain significant variation in brood mass at dispersal.

By generation 43, both parental and larval experimental populations contributed to variation in brood mass at dispersal. No Care broods were still lighter than Full Care broods (linear regression: *F*_1,208_ = 11.733, *p* = 0.001, Figure 1B). However, by this generation the broods raised by No Care parents were also lighter than those cared for by Full Care parents (linear regression: *F*_1,208_ = 6.138, *p* = 0.014), regardless of the larval experimental population (Figure 1B). These effects were greater than, and independent of, the effects of carcass and brood size (Table S1B). Together, these results support Prediction 1 by showing that parental caregiving degraded over time, though more slowly than larval traits for acquiring care.

Prediction 2: If No Care parents had lost the ability to supply care then the duration of their care should be shorter than shown by Full Care parents.

#### (i) Effect of parental sex

There was a significant interaction between the sex of the parent and generation of experimental evolution on the duration of care supplied (ANOVA: *F*_1,734_ = 5.949, *p* = 0.015). In support of Prediction 2, fathers left the brood significantly earlier than mothers in both generations 24 and 43 (ANOVA: *F*_2,736_ = 185.38, *p* = <0.001; Figure 2). Moreover, fathers in generation 24 left their brood earlier than fathers in generation 43 (ANOVA: *F*_1,734_ = 5.949, *p* = 0.015; Figure 2). However, contrary to Prediction 2, there was no significant difference in the duration of care supplied by mothers between the two generations.

**Figure 2.**
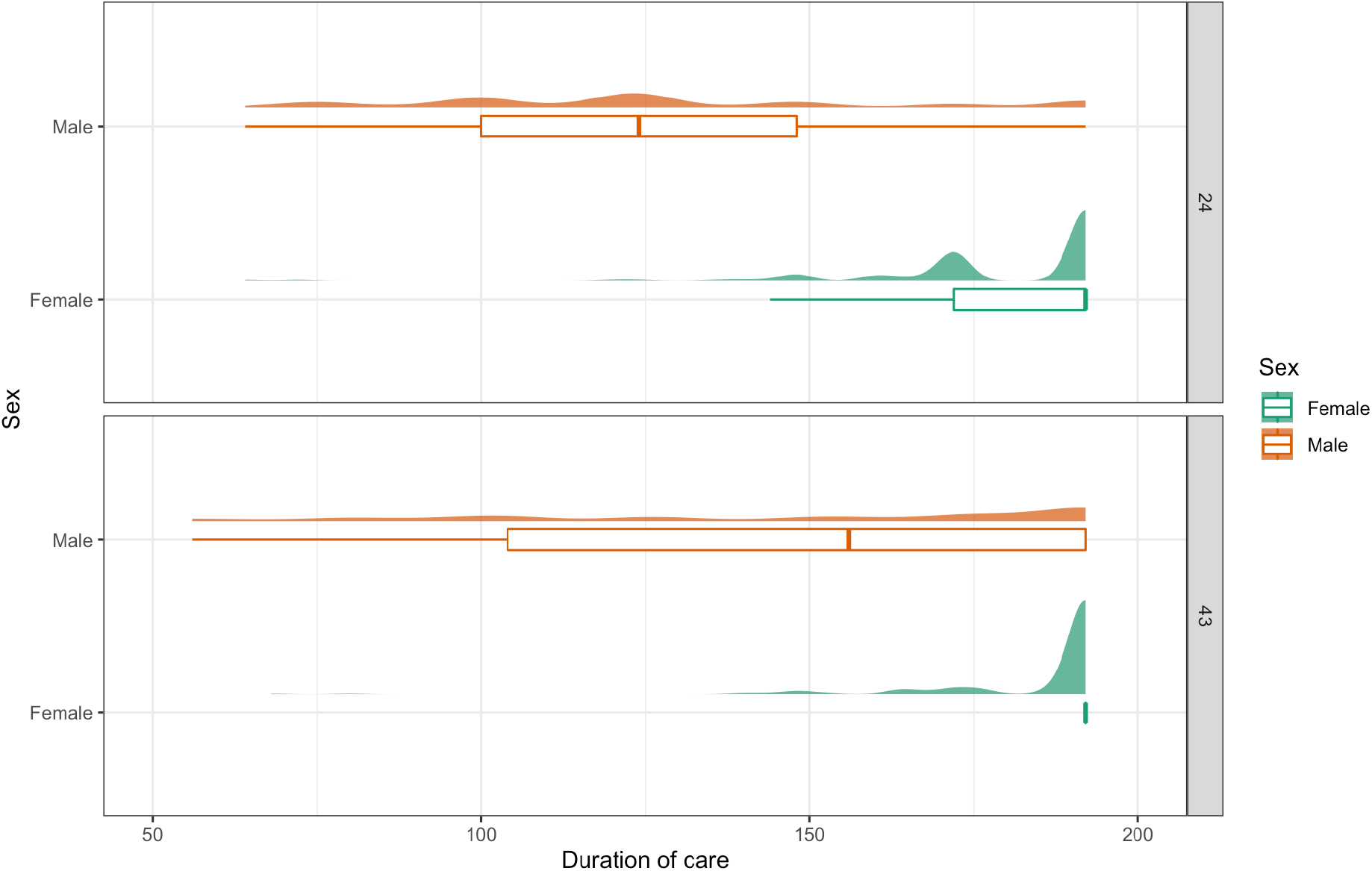
Box and density plots showing the duration of maternal (female, n = 369 individuals) and paternal (male, n = 369 individuals) care in generations 24 and 43. In general, males stayed with the brood for significantly less time than females. Males in generation 24 stayed for less time than males in generation 43. Whiskers extend to the farthest data point which is no more than 1.5 times the interquartile range from the box. Density curves show the distributions of individual points.

We then sought to determine if the duration of parental care influenced average brood mass at dispersal. In generation 24 (Table S1A), the duration of both maternal (female leaving time: linear regression: *F*_1,144_ = 6.393, *p* = 0.013) and paternal care (male leaving time: linear regression: *F*_1,144_ = 6.859, *p* = 0.010) were positively correlated with average brood mass. In generation 43 (Table S2B), only the duration of maternal care contributed to average brood mass (female leaving time: linear regression: *F*_1,208_ = 10.477, *p* = 0.001, male leaving time: linear regression: *F*_1,207_ = 0.004, *p* = 0.950).

#### (ii) Effects of evolutionary lines

We then used a survival analysis at each generation to determine separately how parent and larval experimental populations each independently influenced time of departure from the brood of each parent.

In generation 24, the duration of maternal care was significantly shorter when their brood was drawn from the No Care experimental populations, regardless of the female’s own experimental population of origin (survival model: hazard ratio = 1.749, Wald = 2.085, *p* = 0.028, Figure 3). However, there was no significant effect of parental or larval experimental populations on paternal departure time (Table S2A).

**Figure 3.**
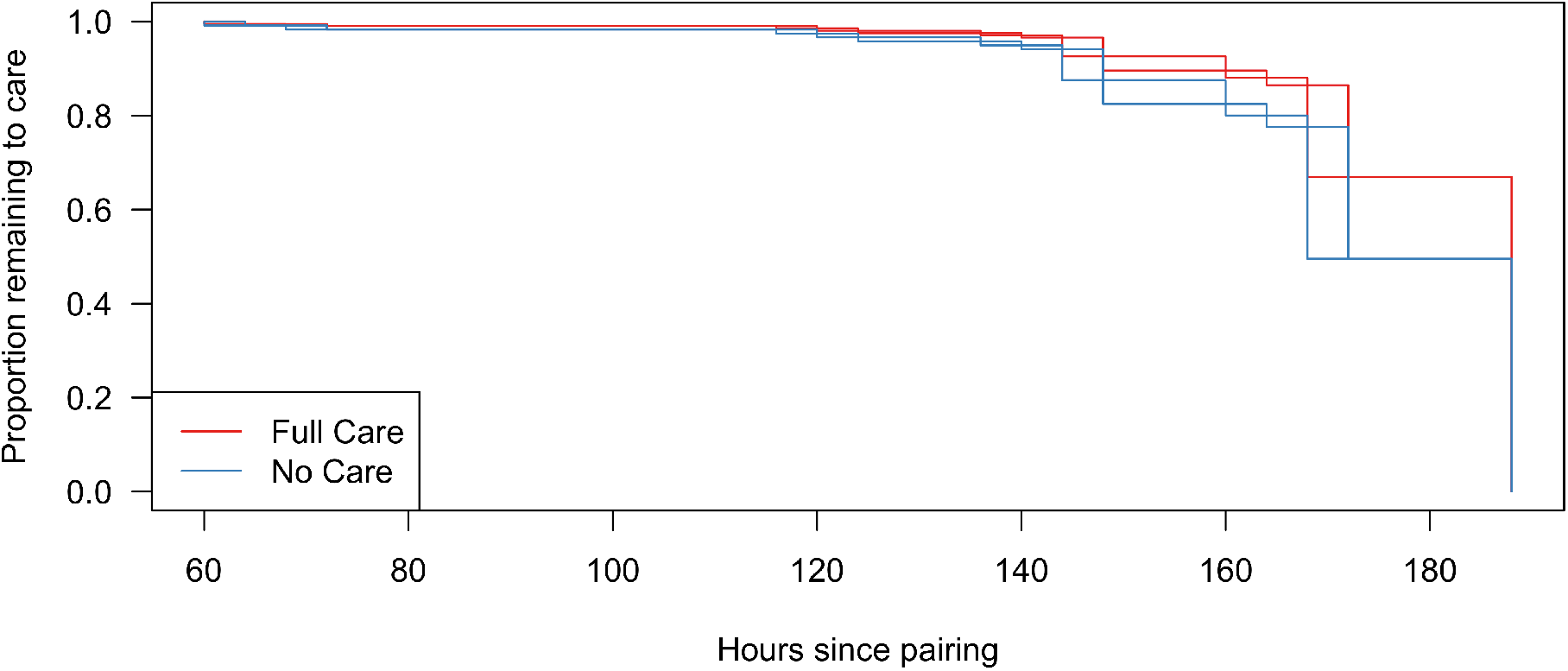
Duration of female care in relation to larval experimental population at generation 24 (Full Care, n = 81; No Care, n = 72 females). The survival curves for female care were significantly affected by experimental population of their current brood (Full Care = red, No Care = blue). Two lines are displayed for each treatment because with interval censored data it is not possible to tell the exact time that the female left, only the time period within which she left. The lines depicted are predicted by the model fitted to the data: for any point between the two lines there is the same likelihood of the parent leaving.

By generation 43 (Table S2B), this pattern had changed. No Care males (survival model: hazard ratio = 1.758, Wald = 3.360, *p* = <0.001, Figure 4) provided care for significantly less time than Full Care males. Although there were no significant effects of experimental population on maternal departure time, the duration of female care was generally shorter when their partner provided less prolonged care (survival model: hazard ratio = 0.993, Wald = −2.027, *p* = 0.021).

**Figure 4.**
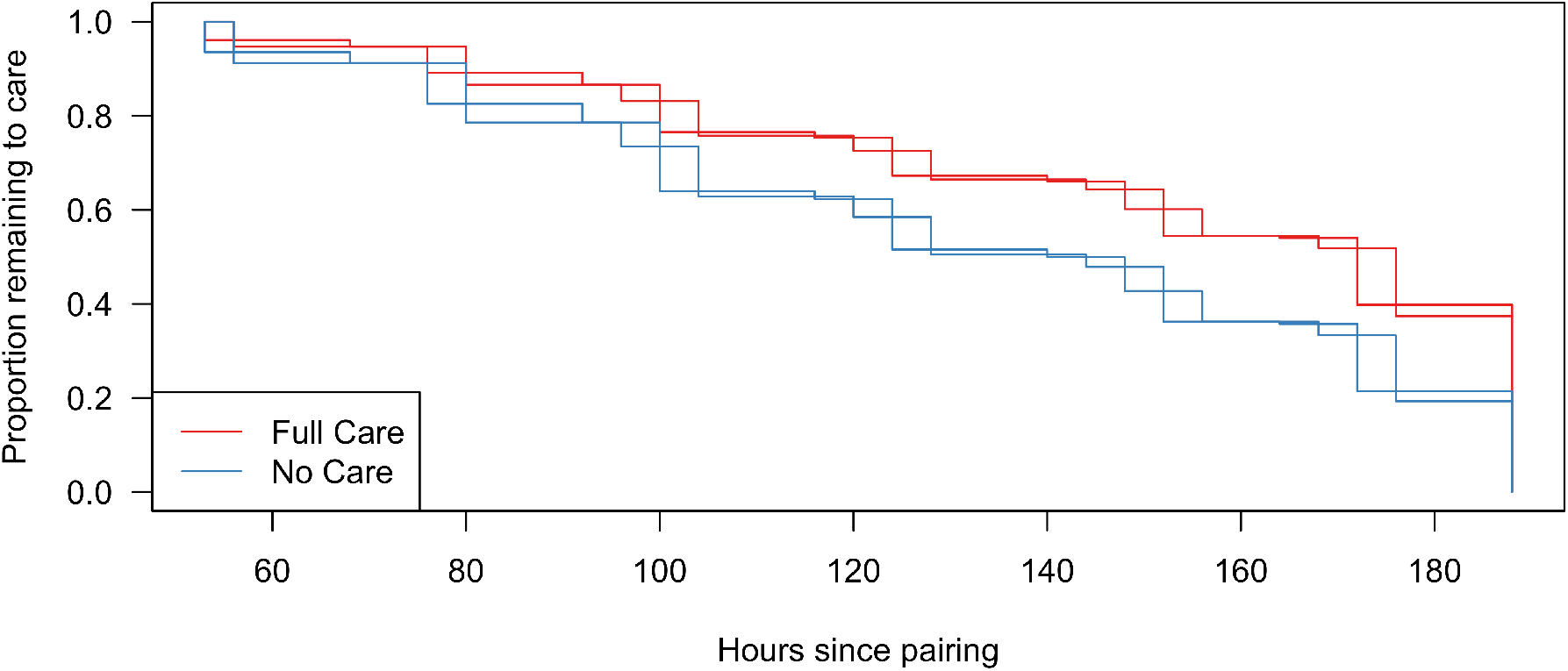
Duration of male care in generation 43 (Full Care, n = 107; No Care, n = 107 males). The survival curves for male care were significantly affected by the male’s experimental population (Full Care = red, No Care = blue), with No Care males caring for less time. Two lines are displayed for each treatment because with interval censored data it is not possible to tell the exact time that the male left, only the time period within which he left. The lines depicted are predicted by the model fitted to the data: for any point between the two lines there is the same likelihood of the parent leaving.

Prediction 3: If No Care larvae had lost the ability to demand care then they should be less inclined to solicit care from parents.

In support of Prediction 3, No Care larvae spent significantly less time begging towards their foster parent than Full Care larvae (linear regression: *F*_1,124_ = 25.042, *p* = <0.001) regardless of the experimental population of their foster parent (Figure 5A). The result was similar when considering total time spent interacting with the parent (linear regression: *F*_1,124_ = 12.979, *p* = 0.001, Figure 5B). No other variables significantly affected larval solicitation behaviours (Table S3).

**Figure 5.**
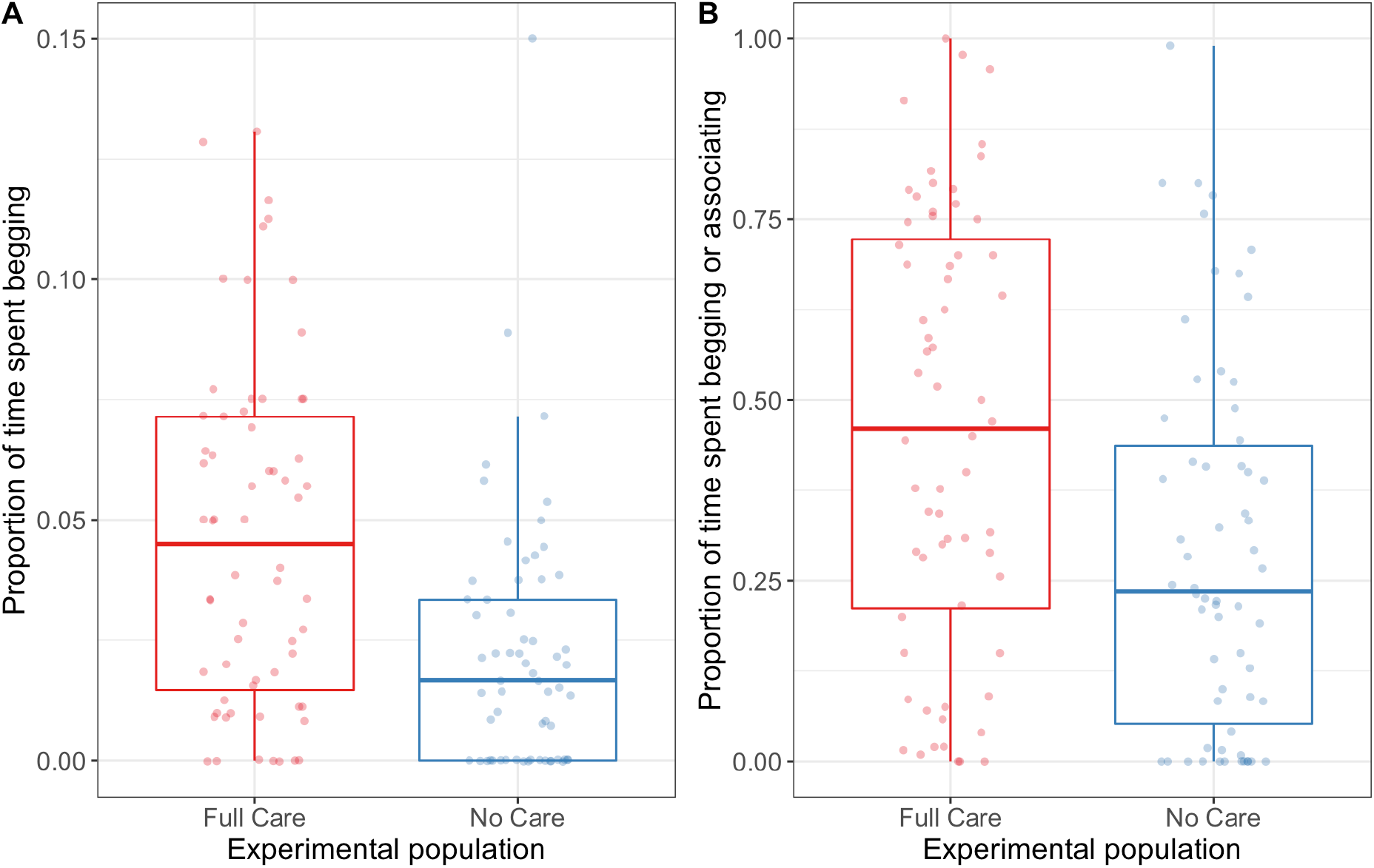
The effect of larval experimental population on A) proportion of time spent begging and B) the proportion of time spent either begging or associating with the parent (n = 126 broods). Whiskers extend to the farthest data point which is no more than 1.5 times the interquartile range from the box. Points show individual broods.

## Discussion

We investigated whether traits connected with offspring demand and parental supply were evolutionarily lost by disrupting this social interaction for generation after generation in experimentally evolving populations. We assessed the extent of loss: 1) indirectly, by partitioning the contribution of the offspring and parental line of origin to average brood mass at dispersal and 2) directly, by quantifying the duration of care supplied by each parent and larval capacity to solicit care. We found support for all three of the predictions we tested.

Our indirect measurements showed that in both generations, No Care larvae (on carrion nests prepared by No Care parents) consistently attained a lighter mass at dispersal than Full Care larvae (on carrion nests prepared by Full Care parents), regardless of whether they were raised by parents from the Full Care or No Care experimental populations. We can think of three possible explanations for these findings, which are not mutually exclusive.

First, offspring from the No Care experimental populations may have developed within smaller eggs, meaning that larvae were smaller from hatching. We do not know whether the eggs produced by the No Care populations were smaller after 24 and 43 generations of experimental evolution, but we have previously shown that development in a No Care environment initially yields smaller adults, and that smaller females lay smaller eggs (Jarrett et al., 2017).

A second explanation is that the carrion nests produced by No Care parents might have provided an inferior environment for offspring development compared with the carrion nests made by Full Care parents. We reject this explanation because we have demonstrated the opposite to be true (Duarte et al., 2021).

The third explanation is that No Care larvae were less effective than Full Care larvae at acquiring resources from parents: the demand for care in the No Care experimental populations was lost because it was not expressed in these populations for so many generations. This interpretation is supported by the direct behavioural measurements, which showed that by generation 48, No Care larvae were less likely to solicit care from parents than Full Care larvae. It also accounts for our findings in generation 24 that all females shortened their duration of care when given No Care larvae to look after. The brood’s failure to solicit care effectively may have contributed to this earlier departure time. On balance, we conclude that the larval capacity to solicit care started to disappear after generations of not being expressed and that this loss might have been evident after just 24 generations (although it is possible that egg size also contributed to a lower average brood mass at dispersal in the No Care larvae, at this point).

Turning to the parents, here we found no evidence for a loss of post-hatching parental behaviour at generation 24, whether we used indirect (brood mass) or direct measurements (departure time). No Care and Full Care parents contributed to a similar extent to average brood mass at dispersal, which was positively correlated with the duration of care supplied by each parent. By generation 43, however, this pattern had changed. Here there was evidence of decline in the No Care male’s supply of post-hatching care, which was now significantly shorter than the care provided by the Full Care males. Furthermore, the duration of male care no longer predicted brood mass at dispersal. In contrast there was no difference between No Care and Full Care females in the time they spent caring for larvae in generation 43, and the duration of maternal care was still positively associated with average brood mass at this point. Nevertheless, shorter periods of male care also curtailed the duration of care supplied by females.

Overall, our experiments show that social behaviours start to disappear when the opportunity to express them is eliminated for multiple generations. Larval demand for care started to disappear first, followed by the provision of post-hatching care by males. Although supply of maternal care decreased in response to weaker demands from larvae, and reduced paternal effort, we found no evidence that No Care mothers had lost any intrinsic capacity to care for larvae - despite being prevented from providing post-hatching care for 43 generations.

Why did we observe contrasting rates of behavioural loss between larvae, males and females? The key to answering this question lies in understanding the mechanisms underpinning these behavioural traits (Heinen-Kay and Zuk, 2019). We can immediately rule out any wider environmental differences, since the experimental populations were kept under identical conditions in the same lab. One possibility is that the genes underpinning each of these traits are simply vulnerable to mutation at different rates or to different extents (Snell-Rood et al., 2009). We have no evidence yet to test whether or not this proposition is true. A different possibility, which is perhaps more likely given the relatively short timescale of this experiment, is that the demand for care and the supply of post-hatching care are each traded against the expression of alternative behavioural traits that offer a different route to gaining fitness (Heinen-Kay and Zuk, 2019). For example, the expression of larval solicitation behaviours could trade-off directly with the expression of self-feeding (Smiseth et al., 2003). In natural populations, there might be standing genetic variation in the threshold at which environmental cues tip the balance between the expression of each larval trait. Selection from the No Care environment could have favoured larvae that were more inclined to express self-feeding at the expense of expressing solicitation behaviours. The recalibration of this trade-off at the population level to favour larval self-feeding in a No Care environment could explain the corresponding loss of larval traits for demanding care.

We can make a similar population-level argument for changes in the expression of male post-hatching care over time. This might be due to a re-balancing of the trade-off with the supply of pre-hatching care and, specifically, the effort devoted to making the carrion nest (de Gasperin et al., 2015). The existence of a trade-off like this is plausible since nest maintenance activity after hatching is known to be negatively genetically correlated with the direct supply of larval care in males (Walling et al., 2008). Furthermore, we have shown elsewhere that by generation 20 the adults from the No Care experimental populations had evolved to put more effort into making the carrion nest than those in the Full Care populations (Duarte et al., 2021) and that this change in nest-making proficiency was due more to males than to females (Jarrett et al., 2022). Additionally, males naturally leave 2 to 5 days earlier than females during the post-hatching period – perhaps because they trade off the potential benefits to their current brood of staying with the potential to sire more offspring away from the carcass (Boncoraglio and Kilner, 2012). This trade-off is stronger for males than females due to the division of labour between the sexes. Male duties of care focus on nest preparation and defence and are completed sooner than female duties of care which focus more on post-hatching care (Walling et al 2008).

Whether an equivalent trade-off exists for females is less clear. The female trait contributing most to fitness before hatching is most likely to be egg size and it is theoretically possible that there is a negative genetic correlation between egg size and the supply of post-hatching care. However, even if such a correlation exists, it is likely to be concealed by a condition-dependent positive phenotypic correlation between egg size and post-hatching care. Females that receive less post-hatching care are smaller and lay correspondingly smaller eggs (Jarrett et al., 2017).

If trade-offs between competing behavioural routes to fitness explain how some traits are lost, then presumably it is the relative strength of selection to recalibrate these trade-offs that explains why rebalancing happened most rapidly in larvae, and then in males, but not in females. Larval mortality in the No Care populations was very high in the first few generations of experimental evolution (Schrader et al., 2017). Any larvae that did not self-feed in the No Care environment would rapidly have been selected against. Selection for recalibrating the male’s trade-off arguably is somewhat weaker, since their nest-building behaviour less directly affects larval survival. The strength of selection on the putative egg-care trade-off in females could be negligible, especially if its expression is masked by female condition.

The general principle emerging from this study is that rapid trait loss is associated with the expression of alternative traits which can enhance fitness more effectively when populations enter a changed environment. This is consistent with previous work analysing trait loss in sexual partners (Heinen-Kay and Zuk, 2019) and with the evolution of trait loss in mutualistic partnerships, where the actions of the partner species compensate for trait loss in the focal species (Ellers et al., 2012). This general principle explains the loss of traits associated with parental care in domesticated bird and mammal species, because breeders often intervene to nurture offspring themselves or to cross-foster them to other strains that provide better offspring care. As a consequence, wild Norway rats have been shown to be more efficient in pup retrieval than domestic mothers (Price and Belanger, 1977), and some strains of domesticated canary (Vriends, 1992) and zebra finch (Blackwell, 1988) are now incapable of raising offspring to independence.

For traits like this, a return to the ancestral environment, even after relatively few generations, is not sufficient to restore ancestral levels of trait expression. There are important implications here for conservation captive breeding programmes, which could inadvertently induce rapid trait loss through any artificial compensatory husbandry techniques that are introduced to promote breeding success in captivity. Interventions like this could prevent the successful reintroduction of species in the wild by causing the loss of key traits needed for survival and reproduction (Araki et al., 2007; Bowkett, 2009; Williams and Hoffman, 2009).

Finally, this study also suggests that although the loss of the interacting phenotype in one individual (e.g. larva, or male) is sufficient to prevent expression of the corresponding phenotype in its social partner (e.g. mother, or female partner), it is not sufficient to cause that trait to decay in the short-term. Nevertheless, unexpressed social traits are likely to be lost in the longer-run, either because of costs to plasticity or because mutations can easily accumulate in traits unseen by natural selection (Snell-Rood et al., 2009, 2018). A complex interplay of neurotransmitter receptor gene expression regulates care in *Nicrophorus vespilloides* (Cunningham et al. 2021) and in future work, it would be interesting to investigate the vulnerability of these genes to mutation.

In general, our study suggests the likelihood of reviving ‘ghosts of adaptation past’ will depend on how long it has been since those adaptive traits were last expressed, and how likely it is that they have been superseded by alternative existing traits that more effectively promote fitness in the new environment. The flexibility of many behavioural traits, and the multiple behavioural routes that are consequently available for achieving similar fitness goals, could mean that some behavioural traits are rapidly substituted and lost following a change in the wider environment.

## Author contributions

RMK and EKB designed the experiments; EKB, SP and NB collected the data. EKB and RM analysed the data. EKB, RM and RMK were involved in writing and editing the manuscript.

## Acknowledgements

This project was supported by a Consolidator’s Grant from the European Research Council (310785 Baldwinian_Beetles), by a Wolfson Merit Award from the Royal Society, The Leverhulme Trust (RPG-2018-232) and The Isaac Newton Trust (18.23(q)), each to RMK. EKB was supported by a Biotechnology and Biological Sciences Research Council PhD studentship (BB/M011194/1). RM was supported by a Biotechnology and Biological Sciences Research Council Future Leaders Fellowship (BB/R01115X/1). NB was supported by an Association for the Study of Animal Behaviour Undergraduate Project Scholarship and by the Balfour-Browne Trust Fund in Cambridge. We thank Chris Swannack and Sue Aspinall for helping with beetle maintainance.

## Supplementary material

**Table S1.** Results of a Gaussian linear model analysing predictors of brood mass in (A) generation 24 and B) generation 43. Significant terms (retained in the minimal model) are shown in bold. All terms included in the maximal model are given and statistics are given for the last model in which the term was retained. “:” represents an interaction between terms.

| Independent variable | Estimate | SE | F | df | p | Estimate | SE | F | df | p |
| --- | --- | --- | --- | --- | --- | --- | --- | --- | --- | --- |
|  | A) Generation 24 |  |  |  |  | B) Generation 43 |  |  |  |  |
| Intercept | -0.945 | 0.268 |  |  |  | -0.695 | 0.307 |  |  |  |
| Carcass mass (g) | **0.134** | **0.022** | **36.811** | **1** | **<0.001** | **0.102** | **0.023** | **19.661** | **1** | **<0.001** |
| Brood size | **0.067** | **0.003** | **540.18** | **1** | **<0.001** | **-0.080** | **0.003** | **748.16** | **1** | **<0.001** |
| Current parents' experimental population (No Care) | -0.049 | 0.037 | 1.767 | 1 | 0.186 | **-0.093** | **0.036** | **6.775** | **1** | **0.010** |
| Larval experimental population (No Care) | **-0.168** | **0.037** | **20.274** | **1** | **<0.001** | **-0.131** | **0.035** | **13.744** | **1** | **<0.001** |
| Transferred? (Yes) | 0.026 | 0.038 | 0.463 | 1 | 0.497 | **-0.074** | **0.036** | **4.358** | **1** | **0.038** |
| Male duration of care | **0.001** | **0.001** | **4.921** | **1** | **0.028** | <0.001 | <0.001 | 0.173 | 1 | 0.678 |
| Female duration of care | **0.002** | **0.001** | **6.005** | **1** | **0.016** | **0.003** | **0.001** | **8.565** | **1** | **0.004** |
| Block (2) | **-0.114** | **0.038** | **8.885** | **1** | **0.003** | **-0.070** | **0.033** | **4.488** | **1** | **0.035** |
| Current parents' line:Larval experimental population | 0.059 | 0.088 | 0.448 | 1 | 0.505 | -0.066 | 0.081 | 0.659 | 1 | 0.418 |

**Table S2.** Results of semi-parametric Cox’s proportional models of parental leaving times in A) generation 24 and B) generation 43. Terms retained in the minimal model are shown in bold. All terms included in the maximal model are given and statistics given are for the last model in which the term was retained “:” represents an interaction between terms.

| Parent | Independent Variable | Parameter Estimate | Hazard Ratio | SE | z | p | Parameter Estimate | Hazard Ratio | SE | z | p |
| --- | --- | --- | --- | --- | --- | --- | --- | --- | --- | --- | --- |
|  |  | **A) Generation 24** |  |  |  |  | **B) Generation 43** |  |  |  |  |
| Male | Male evolutionary population (No Care) | 0.073 | 1.076 | 0.170 | 0.430 | 0.680 | **0.564** | **1.758** | **0.168** | **3.360** | **<0.001** |
|  | Brood evolutionary population (No Care) | 0.022 | 1.022 | 0.218 | 0.099 | 0.908 | -0.322 | 0.725 | 0.177 | -1.823 | 0.062 |
|  | Carcass mass (g) | 0.011 | 1.011 | 0.013 | 0.081 | 0.927 | 0.106 | 1.112 | 0.105 | 1.009 | 0.317 |
|  | Transferred? (Yes) | 0.151 | 1.163 | 0.169 | 0.893 | 0.402 | -0.116 | 0.891 | 0.167 | -0.691 | 0.505 |
|  | Brood size | 0.003 | 1.003 | 0.013 | 0.224 | 0.833 | -0.007 | 0.993 | 0.017 | -0.380 | 0.659 |
|  | Block (1) | 0.155 | 1.168 | 0.197 | 0.791 | 0.382 | **0.365** | **1.441** | **0.169** | **2.160** | **0.026** |
| Female | Female evolutionary population (No Care) | 0.548 | 1.730 | 0.284 | 1.934 | 0.426 | 0.082 | 1.086 | 0.335 | 0.246 | 0.794 |
|  | Brood evolutionary population (No Care) | **0.559** | **1.749** | **0.268** | **2.085** | **0.028** | -0.132 | 0.876 | 0.266 | -0.497 | 0.639 |
|  | Carcass mass (g) | -0.071 | 0.932 | 0.151 | -0.469 | 0.673 | -0.170 | 0.843 | 0.188 | -0.904 | 0.390 |
|  | Transferred? (Yes) | 0.020 | 1.020 | 0.389 | 0.052 | 0.946 | 0.027 | 1.027 | 0.350 | 0.077 | 0.929 |
|  | Brood size | 0.038 | 1.038 | 0.022 | 1.675 | 0.061 | 0.024 | 1.024 | 0.024 | 0.986 | 0.318 |
|  | Block (1) | 0.254 | 1.289 | 0.230 | 0.849 | 0.355 | 0.096 | 1.100 | 0.295 | 0.324 | 0.736 |
|  | Male duration of care | -0.007 | 0.993 | 0.004 | -1.671 | 0.057 | **-0.007** | **0.993** | **0.004** | **-2.027** | **0.021** |

**Table S3.**
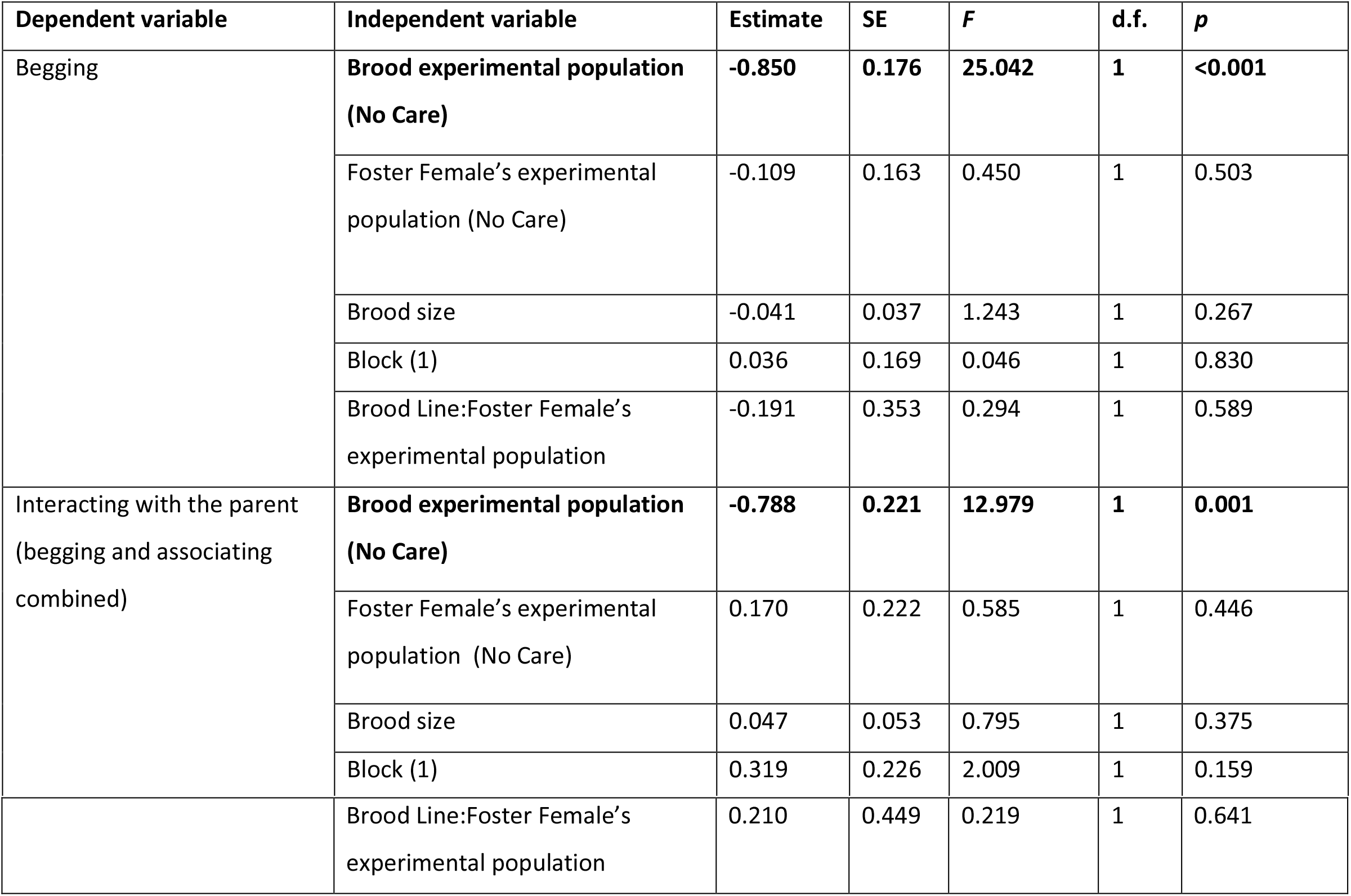
Quasi-binomial GLMs fitted to explain variation in the proportion of time spent begging and the proportion of time spent either begging or associating with the parent. ‘Begging’ was defined as occurring when a larva reached up to touch the parents with its forelegs and ‘associating’ was defined as a larva being within a pronotum’s width of the parent’s body. All terms included in the maximal model are shown, as well as their contribution to the final model. Terms retained in the minimal model are shown in bold. “:” represents an interaction between terms.

| Dependent variable | Independent variable | Estimate | SE | F | d.f. | p |
| --- | --- | --- | --- | --- | --- | --- |
| Begging | **Brood experimental population (No Care)** | **-0.850** | **0.176** | **25.042** | **1** | **<0.001** |
|  | Foster Female’s experimental population (No Care) | -0.109 | 0.163 | 0.450 | 1 | 0.503 |
|  | Brood size | -0.041 | 0.037 | 1.243 | 1 | 0.267 |
|  | Block (1) | 0.036 | 0.169 | 0.046 | 1 | 0.830 |
|  | Brood Line:Foster Female’s experimental population | -0.191 | 0.353 | 0.294 | 1 | 0.589 |
| Interacting with the parent (begging and associating combined) | **Brood experimental population (No Care)** | **-0.788** | **0.221** | **12.979** | **1** | **0.001** |
|  | Foster Female’s experimental population (No Care) | 0.170 | 0.222 | 0.585 | 1 | 0.446 |
|  | Brood size | 0.047 | 0.053 | 0.795 | 1 | 0.375 |
|  | Block (1) | 0.319 | 0.226 | 2.009 | 1 | 0.159 |
|  | Brood Line:Foster Female's<br>experimental population | 0.210 | 0.449 | 0.219 | 1 | 0.641 |

**Figure S1:**
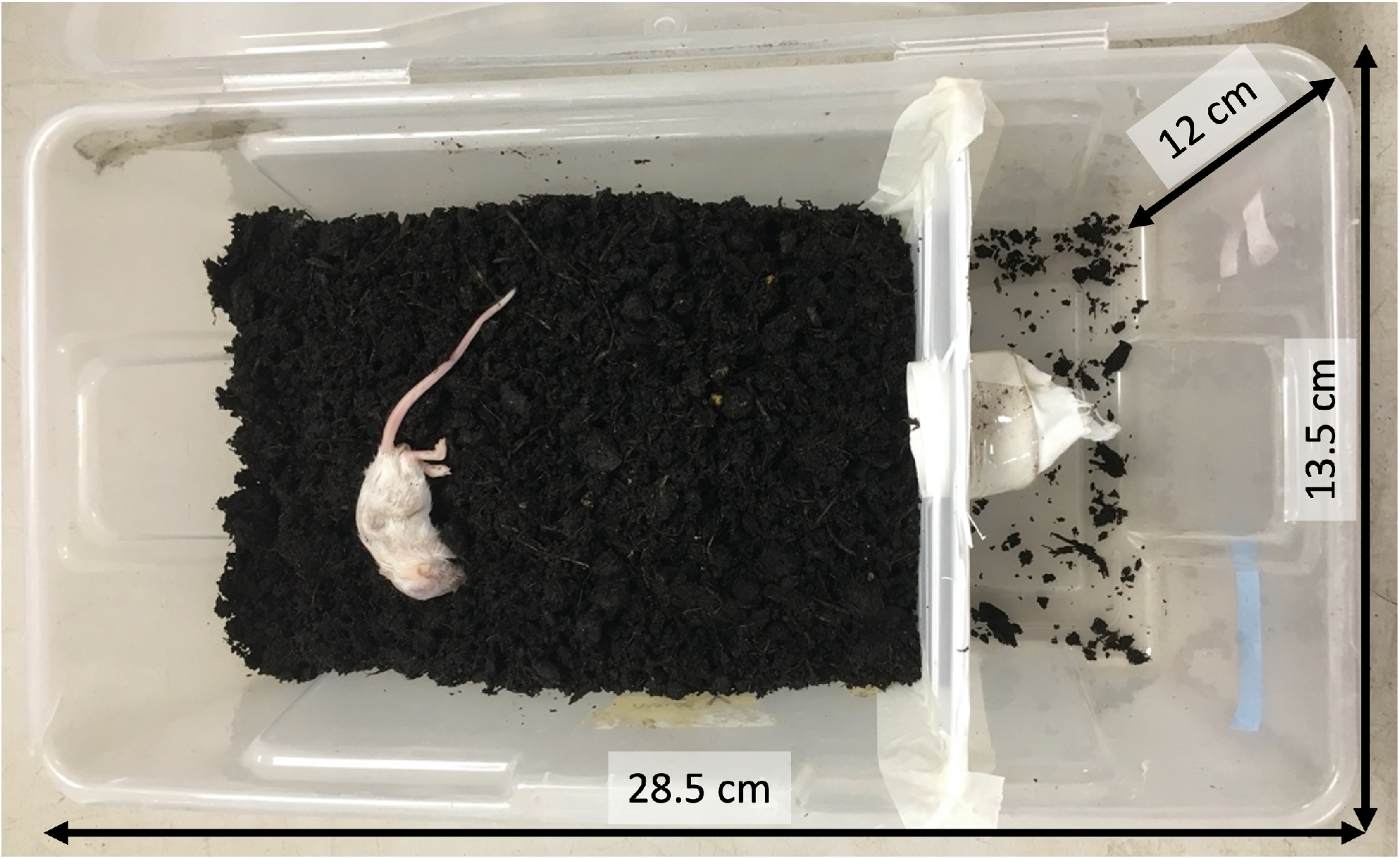
The breeding boxes used in Experiment 1 with breeding compartment and “escape chamber”.

**Figure S2:**
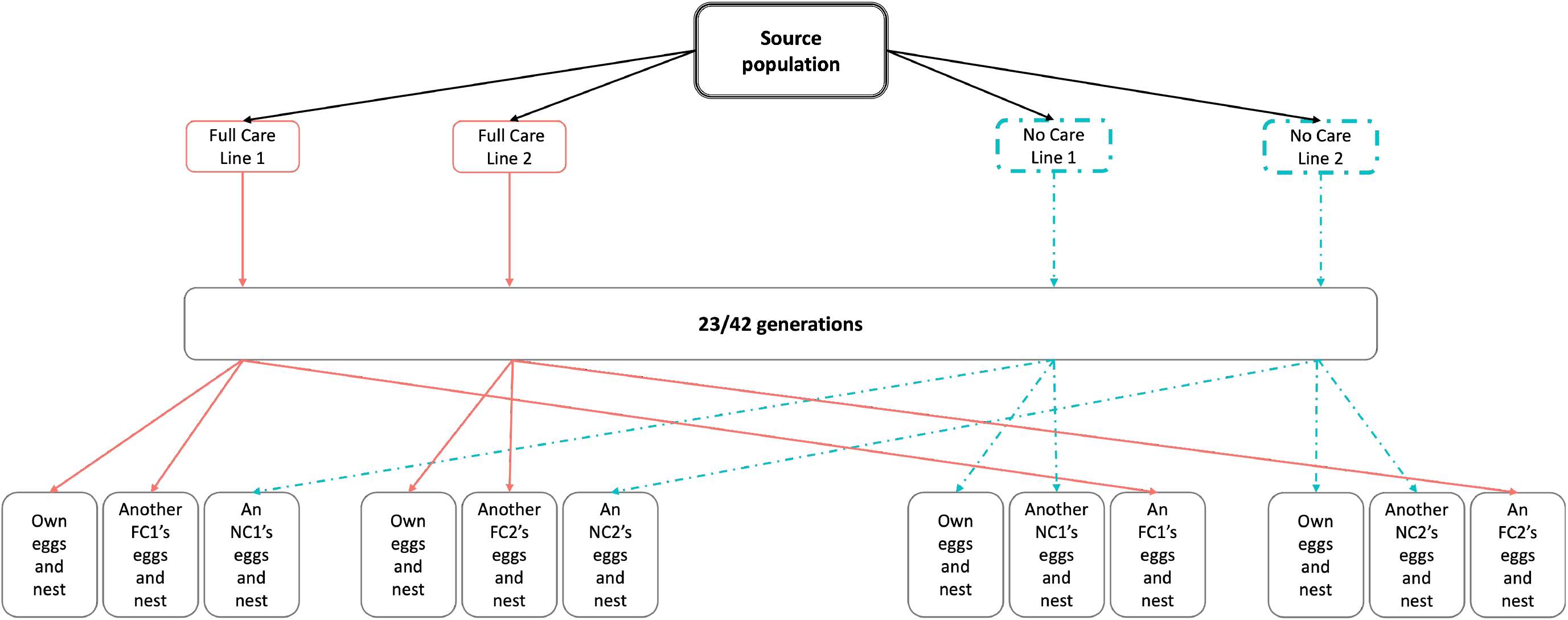
The design of the experimental evolving populations and Experiment 1

## Notes

### Competing Interest Statement

The authors have declared no competing interest.

### Summary of Updates

This manuscript has been updated following a first round of peer review. The major points of revision are an improvement in the presentation of figures for clarity, the removal of one analysis which was considered unnecessary, and the use of total brood mass instead of average larva mass as the dependent variable in applicable analyses.

## References

Anderson-Bergman, C. (2017). icenReg: regression models for interval censored data in R. Journal of Statistical Software 81, 1–23. https://doi.org/10.18637/jss.v081.i12.

Araki, H., Cooper, B., and Blouin, M.S. (2007). Genetic Effects of Captive Breeding Cause a Rapid, Cumulative Fitness Decline in the Wild. Science 318, 100. https://doi.org/10.1126/science.1145621.

Bailey, N.W., and Moore, A.J. (2018). Evolutionary Consequences of Social Isolation. Trends in Ecology & Evolution 33, 595–607. https://doi.org/10.1016/j.tree.2018.05.008.

Blackwell, C. (1988). Keeping and Breeding Zebra Finches: The Complete Type Standard Guide (Blandford Press).

Bladon, E.K., English, S., Pascoal, S., and Kilner, R.M. (2020). Early-life effects on body size in each sex interact to determine reproductive success in the burying beetle Nicrophorus vespilloides. Journal of Evolutionary Biology 33, 1725–1734. https://doi.org/10.1111/jeb.13711.

Boncoraglio, G., and Kilner, R.M. (2012). Female burying beetles benefit from male desertion: sexual conflict and counter-adaptation over parental investment. PloS One 7,e31713. https://doi.org/10.1371/journal.pone.0031713.

Bowkett, A.E. (2009). Recent Captive-Breeding Proposals and the Return of the Ark Concept to Global Species Conservation. Conservation Biology 23, 773–776. https://doi.org/10.1111/j.1523-1739.2008.01157.x.

Cotter, S., and Kilner, R.M. (2010). Sexual division of antibacterial resource defence in breeding burying beetles, *Nicrophorus vespilloides*. Journal of Animal Ecology 79, 35–43. https://doi.org/10.1111/j.1365-2656.2009.01593.x.

Crawley, M.J. (2007). Multiple regression. In The R Book, (Chichester: John Wiley & Sons, Ltd), pp. 569–591.

Cunningham, C.B., Khana, D., Carter, A., McKinney, E.C., and Moore, A.J. (2021). Survey of neurotransmitter receptor gene expression into and out of parental care in the burying beetle Nicrophorus vespilloides. Ecology and Evolution 11, 14282–14292. https://doi.org/10.1002/ece3.8144.

Davies, N.B. (2000). Cuckoos, Cowbirds and Other Cheats (London: Poyser).

De Gasperin, O., Duarte, A., and Kilner, R.M. (2015). Interspecific interactions explain variation in the duration of paternal care in the burying beetle. Animal Behaviour 109, 199–207. https://doi.org/10.1016/j.anbehav.2015.08.014.

DeWitt, T.J., Sih, A., and Wilson, D.S. (1998). Costs and limits of phenotypic plasticity. Trends in Ecology and Evolution 13. https://doi.org/10.1016/S0169-5347(97)01274-3.

Duarte, A., Rebar, D., Hallett, A.C., Jarrett, B.J.M., and Kilner, R.M. (2021). Evolutionary change in the construction of the nursery environment when parents are prevented from caring for their young directly. Proceedings of the National Academy of Sciences 118,e2102450118. https://doi.org/10.1073/pnas.2102450118.

Eggert, A.-K., Reinking, M., and Müller, J.K. (1998). Parental care improves offspring survival and growth in burying beetles. Animal Behaviour 55, 97–107. https://doi.org/10.1006/anbe.1997.0588.

Ellers, J., Kiers, E.T., Currie, C.R., McDonald, B.R., and Visser, B. (2012). Ecological interactions drive evolutionary loss of traits. Ecology Letters 15, 1071–1082. https://doi.org/10.1111/j.1461-0248.2012.01830.x.

Heinen-Kay, J.L., and Zuk, M. (2019). When does sexual signal exploitation lead to signal loss? Frontiers in Ecology and Evolution 7. https://doi.org/10.3389/fevo.2019.00255.

Jarrett, B.J.M., Schrader, M., Rebar, D., Houslay, T.M., and Kilner, R.M. (2017). Cooperative interactions within the family enhance the capacity for evolutionary change in body size. Nature Ecology & Evolution 1, 0178. http://dx.doi.org/10.1038/s41559-017-0.

Jarrett, B.J.M., Evans, E., Haynes, H.B., Leaf, M.R., Rebar, D., Duarte, A., Schrader, M., and Kilner, R.M. (2018a). A sustained change in the supply of parental care causes adaptive evolution of offspring morphology. Nature Communications 9, 3987. https://doi.org/10.1038/s41467-018-06513-6.

Jarrett, B.J.M., Rebar, D., Haynes, H.B., Leaf, M.R., Halliwell, C., Kemp, R., and Kilner, R.M. (2018b). Adaptive evolution of synchronous egg-hatching in compensation for the loss of parental care. Proceedings of the Royal Society B: Biological Sciences 285, 20181452. https://doi.org/10.1098/rspb.2018.1452.

Jarrett, B.J.M., Mashoodh, R., Issar, S., Pascoal, S., Rebar, D., Sun, S.-J., Schrader, M., and Kilner, R.M. (2022). Multilevel selection leads to divergent coadaptation of care-giving parents during pre-hatching parental care. BioRxiv 2022.05.23.493134. https://doi.org/10.1101/2022.05.23.493134.

Kalinová, B., Podskalská, H., Růžička, J., and Hoskovec, M. (2009). Irresistible bouquet of death—how are burying beetles (Coleoptera: Silphidae: *Nicrophorus)* attracted by carcasses. Naturwissenschaften 96, 889–899. https://doi.org/10.1007/s00114-009-0545-6.

Lahti, D.C. (2006). Persistence of egg recognition in the absense of cuckoo brood parasitism: pattern and mechanism. Evolution 60, 157–168. https://doi.org/10.1111/j.0014-3820.2006.tb01090.x.

Lahti, D.C., Johnson, N.A., Ajie, B.C., Otto, S.P., Hendry, A.P., Blumstein, D.T., Coss, R.G., Donohue, K., and Foster, S.A. (2009). Relaxed selection in the wild. Trends in Ecology & Evolution 24, 487–496. https://doi.org/10.1016/j.tree.2009.03.010.

Mäenpää, M.I., and Smiseth, P.T. (2017). Egg size, begging behaviour and offspring fitness in Nicrophorus vespilloides. Animal Behaviour 134, 201–208. https://doi.org/10.1016/j.anbehav.2017.10.014.

McGlothlin, J.W., Moore, A.J., Wolf, J.B., and Brodie III, E.D. (2010). Interacting phenotypes and the evolutionary process. III. Social evolution. Evolution 64, 2558–2574. https://doi.org/10.1111/j.1558-5646.2010.01012.x.

Moore, A.J., Brodie III, E.D., and Wolf, J.B. (1997). Interacting phenotypes and the evolutionary process: I. Direct and indirect genetic effects of social interactions. Evolution 51, 1352–1362. https://doi.org/10.1111/j.1558-5646.1997.tb01458.x.

Müller, J.K., and Eggert, A.-K. (1989). Paternity assurance by “helpful” males: adaptations to sperm competition in burying beetles. Behavioral Ecology and Sociobiology 24, 245–249. https://doi.org/10.1007/BF00295204.

Müller, J.K., and Eggert, A.-K. (1990). Time-dependent shifts between infanticidal and parental behavior in female burying beetles a mechanism of indirect mother-offspring recognition. Behavioral Ecology and Sociobiology 27, 11–16. https://doi.org/10.1007/BF00183307.

Pascoal, S., Jarrett, B., Evans, E., and Kilner, R. (2018). Superior stimulation of female fecundity by subordinate males provides a mechanism for telegony. Evolution Letters 2, 114–125. https://doi.org/10.1002/evl3.45.

Pigliucci, M., Murren, C.J., and Schlichting, C.D. (2006). Phenotypic plasticity and evolution by genetic assimilation. Journal of Experimental Biology 209, 2362–2367. https://doi.org/10.1242/jeb.02070.

Price, E.O., and Belanger, P.L. (1977). Maternal behavior of wild and domestic stocks of Norway rats. Behavioral Biology 20, 60–69. https://doi.org/10.1016/S0091-6773(77)90511-9.

R Core Team (2019). R: A language and environment for statistical computing (Vienna, Austria: R Foundation for Statistical Computing).

Rayner, J.G., Sturiale, S.L., and Bailey, N.W. (2022). The persistence and evolutionary consequences of vestigial behaviours. Biological Reviews 97, 1389–1407. https://doi.org/10.1111/brv.12847.

Rebar, D., Bailey, N.W., Jarrett, B.J.M., and Kilner, R.M. (2020). An evolutionary switch from sibling rivalry to sibling cooperation, caused by a sustained loss of parental care. Proceedings of the National Academy of Sciences 201911677. https://doi.org/10.1073/pnas.1911677117.

Robert, M., and Sorci, G. (1999). Rapid increase of host defence against brood parasites in a recently parasitized area: the case of village weavers in Hispaniola. Proceedings of the Royal Society B: Biological Sciences 266, 941–946. https://doi.org/10.1098/rspb.1999.0727.

Robinson, B.W., and Dukas, R. (1999). The Influence of Phenotypic Modifications on Evolution: The Baldwin Effect and Modern Perspectives. Oikos 85, 582–589. https://doi.org/10.2307/3546709.

Rothstein, S.I. (2001). Relic behaviours, coevolution and the retention versus loss of host defences after episodes of avian brood parasitism. Animal Behaviour 61, 95–107. https://doi.org/10.1006/anbe.2000.1570.

Scheiner, S.M., and Levis, N.A. (2021). The Loss of Phenotypic Plasticity Via Natural Selection: Genetic Assimilation. In Phenotypic Plasticity & Evolution: Causes, Consequences, Controversies, (Boca Raton: CRC Press), pp. 161–177.

Schrader, M., Jarrett, B.J.M., and Kilner, R.M. (2015a). Parental care masks a density-dependent shift from cooperation to competition among burying beetle larvae. Evolution; International Journal of Organic Evolution 69, 1077–1084. https://doi.org/10.1111/evo.12615.

Schrader, M., Jarrett, B.J.M., and Kilner, R.M. (2015b). Using Experimental Evolution to Study Adaptations for Life within the Family. The American Naturalist 185, 610–619. https://doi.org/10.1086/680500.

Schrader, M., Jarrett, B.J.M., Rebar, D., and Kilner, R.M. (2017). Adaptation to a novel family environment involves both apparent and cryptic phenotypic changes. Proceedings of the Royal Society B: Biological Sciences 284, 20171295. https://doi.org/10.1098/rspb.2017.1295.

Smiseth, P.T., and Parker, H.J. (2008). Is there a cost to larval begging in the burying beetle Nicrophorus vespilloides? Behavioral Ecology 19, 1111–1115. https://doi.org/10.1093/beheco/arn101.

Smiseth, P.T., Darwell, C.T., and Moore, A.J. (2003). Partial begging: an empirical model for the early evolution of offspring signalling. Proceedings of the Royal Society B: Biological Sciences 270, 1773–1777. https://doi.org/10.1098/rspb.2003.2444.

Smiseth, P.T., Andrews, C., Brown, E., and Prentice, P.M. (2010). Chemical stimuli from parents trigger larval begging in burying beetles. Behavioral Ecology 21, 526–531. https://doi.org/10.1093/beheco/arq019.

Snell-Rood, E.C., Van Dyken, J.D., Cruickshank, T., Wade, M.J., and Moczek, A.P. (2009). Toward a population genetic framework of developmental evolution: the costs, limits, and consequences of phenotypic plasticity. BioEssays 32, 71–81. https://doi.org/10.1002/bies.200900132.

Snell-Rood, E.C., Kobiela, Megan E., Sikkink, Kristin L., and Shephard, A.M. (2018). Mechanisms of Plastic Rescue in Novel Environments. Annual Review of Ecology, Evolution, and Systematics 49, 331–354. https://doi.org/10.1146/annurev-ecolsys-110617-062622.

Vriends, M.M. (1992). The New Canary Handbook (Barron’s Educational Series Inc.,U.S.).

Waddington, C.H. (1953). Genetic assimilation of an acquired character. Evolution 7, 118–126. https://doi.org/10.1111/j.1558-5646.1953.tb00070.x.

Walling, C.A., Stamper, C.E., Smiseth, P.T., and Moore, A.J. (2008). The quantitative genetics of sex differences in parenting. Proceedings of the National Academy of Sciences 105, 18430. https://doi.org/10.1073/pnas.0803146105.

Wickham, H., Averick, M., Bryan, J., Chang, W., D’Agostino McGowan, L., Romain, F., Grolemund, G., Hayes, A., Henry, L., Hester, J., et al. (2019). Welcome to the tidyverse. Journal of Open Source Software 4, 1686. https://doi.org/10.21105/joss.01686.

Williams, S.E., and Hoffman, E.A. (2009). Minimizing genetic adaptation in captive breeding programs: A review. Biological Conservation 142, 2388–2400. https://doi.org/10.1016/j.biocon.2009.05.034.

Wolf, J.B., Brodie III, E.D., Cheverud, J.M., Moore, A.J., and Wade, M.J. (1998). Evolutionary consequences of indirect genetic effects. Trends in Ecology & Evolution 13, 64–69. https://doi.org/10.1016/S0169-5347(97)01233-0.

